# Reprogramming progressive cells display low CAG promoter activity

**DOI:** 10.1101/2020.03.03.975664

**Authors:** Xiao Hu, Qiao Wu, Jian Zhang, Xinyue Chen, Amaleah Hartman, Anna Eastman, Shangqin Guo

## Abstract

There is wide variability in the propensity of somatic cells to reprogram into pluripotency in response to the Yamanaka factors. How to segregate these variability to enrich for cells of specific traits that reprogram efficiently remains challenging. Here we report that the variability in reprogramming propensity is associated with the activity of the MKL1/SRF transcription factor and concurs with small cell size as well as rapid cell cycle. Reprogramming progressive cells can be prospectively identified by their low activity of a widely used synthetic promoter, CAG. CAG_low_ cells arise and expand during cell cycle acceleration in the early reprogramming culture of both mouse and human fibroblasts. Our work illustrate a molecular scenario underlying the distinct reprogramming propensities and demonstrate a convenient practical approach for their enrichment.

## Introduction

The somatic cells amenable to switching into pluripotency upon expression of the Yamanaka factors are considered to exist stochastically (Buganim et al., 2012; Hanna et al., 2009; Yamanaka, 2009). Within such a model, the rare cells known to have high reprogramming potential, such as the privileged cells or the poised/elite/winner cells (Di Stefano et al., 2016), could represent extreme cellular states existing by chance. With the advent of various single cell genomics, it is now possible to define the expression of large numbers of genes in many individual cells (Battich et al., 2015; Buettner et al., 2015;Cote et al., 2016; Klein et al., 2015; Newman et al., 2006), capturing data sufficient for quantitative assessment of the stochastic nature in gene expression of single cells. Surprisingly, such studies revealed that the expression level of most genes is minimally stochastic, and can in fact be reliably predicted (Battich et al., 2015). Intriguingly, the most predicative parameters of gene expression heterogeneity are DNA content, indicative of cell cycle status, and cell size/shape. These findings suggest the possibility that if the rare cells of certain cell cycle behavior and/or cell size/shape can be prospectively identified or enriched, the stochasticity of somatic cell progressing into pluripotency may be minimized.

Cell size and shape is largely and collectively controlled by the concentration and conformation of cytoskeletal proteins. The expression of many cytoskeletal genes is under the control of MKL1 (Megakaryoblast Leukemia 1)/SRF (serum response factor), via the consensus CArG motif in their promoters. Important MKL1/SRF target genes include many members of the actin genes, and *SRF* itself (Spencer and Misra, 1996). Besides cytoskeletal genes, another major class of SRF target genes are the “immediately early genes”, such as the AP1 family of transcription factors *c-jun*, *c-fos and Fra1* (Bahrami and Drablos, 2016; Cen et al., 2004; Lee et al., 2010; Treisman, 1990). These transcription factors are now known to antagonize Yamanka reprogramming, or established pluripotency by redefining the enhancer-Pol II relationship (Chronis et al.,2017; Hamilton et al., 2019; Liu et al., 2015). We have recently reported that MKL1/SRF activity potently prevents the activation of mature pluripotency by hindering chromatin accessibility via an actin-LINC-dependent process (Hu et al., 2019). Taken together, these evidences suggest that the transcriptional activity of the ubiquitously expressed SRF could potentially serve as an indicator for how likely a somatic cell could progress into pluripotency.

In this report, we reveal that the fast cycling cells are small in size and display reduced SRF target gene expression. Low endogenous SRF activity can be conveniently captured by a synthetic promoter, the CAG promoter which derives part of its sequence from the chicken actin promoter/enhancer (Ng et al., 1989; Stoflet et al., 1992). Prospective isolation of CAG_low_ cells significantly enriched for reprogramming progressive cells in both mouse and human fibroblasts. Our work demonstrates that the inherent difference in reprogramming potential can be purified, at least partially, by the activity of a ubiquitously expressed transcription factor SRF.

## Results

### Cells expressing low CAG promoter activity emerge during early reprogramming

Given that SRF target genes, the cytoskeletal genes and immediate early genes, both antagonize pluripotency, we wondered whether cells with diminished SRF activity represent the cells of higher reprogramming potential. SRF drives expression of target genes by binding to the consensus CArG sequences in their promoters, thus transgenes under the control of functional CArG elements might provide a convenient measure of SRF activity. Since the key functional element of the widely used CAG promoter contains two well conserved CArG sites (Ng et al., 1989; Stoflet et al., 1992), we tested the intensity of fluorescent reporters driven by the CAG promoter during early reprogramming.

We first assessed reprogramming of mouse embryonic fibroblasts (MEFs) derived from a transgenic mouse line expressing an H2B-GFP fusion protein driven by the CAG promoter (Hadjantonakis and Papaioannou, 2004), crossed with the *Rosa26:rtTA* allele (Hochedlinger et al., 2005). A lentiviral vector directing doxycycline (Dox) inducible OSKM polycistronic cassette (Carey et al., 2009) that also includes mCherry was used to drive Yamanaka factor expression (Figure 1A). With this system, reprogramming cells are contained within the mCherry^+^ cells after Dox is added. Cells with lower SRF activity should display lower CAG:H2B-GFP fluorescence intensity.

**Figure 1.**
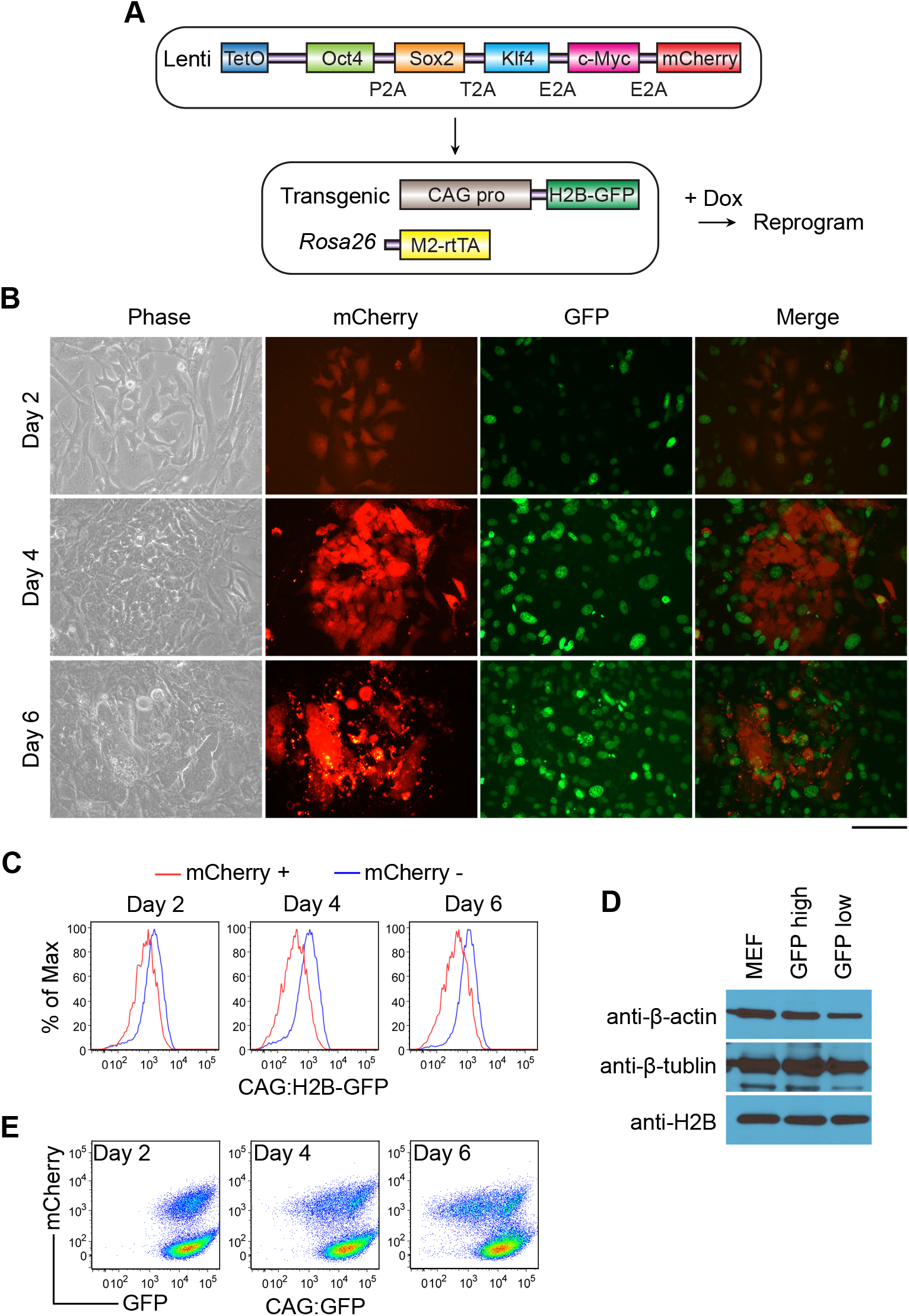
Reprogramming progressive cells show reduced CAG promoter activity. (A) Schematic representation of the experimental design of reprogramming. (B) Downregulation of CAG:H2B-GFP in reprogramming cells as revealed by fluorescence microscopy. mCherry marks OSKM-expressing cells. Scale bar: 100 μm. (C) FACS analysis of the CAG:H2B-GFP MEFs undergoing reprogramming. OSKMmCherry^+^ cells display reduced GFP signal. (D) Western blot of whole cell lysates from bulk MEFs or subsets isolated from reprogramming cultures based on H2B-GFP intensity. With β-tubulin as loading control, the protein levels relative to MEFs are 0.94 and 1.02 for endogenous H2B, 0.70 and 0.53 for β-actin in H2B-GFP_high_ and H2B-GFP_low_ cells, respectively. (E) FACS analysis of the CAG:GFP MEF cells undergoing reprogramming. A fraction of OSKMmCherry+ cells decreased GFP signal as reprogramming continued.

We examined whether CAG:H2B-GFP intensity decreases during early reprogramming to reflect the decreasing MKL1/SRF activity (Hu et al., 2019). Shortly after Dox addition, there was a marked down-regulation of CAG:H2B-GFP intensity within the mCherry^+^ population (Figures 1B-C), consistent with reduced activity of MKL1/SRF. The decreased H2B-GFP signal was not a result from down-regulation of the H2B moiety, because endogenous H2B protein did not decrease in the CAG:H2B-GFP_low_ cells (Figure 1D). To rule out the possibility that the down-regulation of the CAG:H2B-GFP signal was mediated by the particular transgene integration site, we reprogramming MEFs derived from another independent transgenic mouse line (Okabe et al., 1997) similarly crossed with the *Rosa26:rtTA* allele (Hochedlinger et al., 2005). These MEFs express GFP without H2B under the same CAG promoter but presumably have a different transgene integration site. Similar to the observations with CAG:H2B-GFP MEFs, the CAG:GFP level was significantly reduced in the mCherry^+^ cells early in reprogramming (Figure 1E). Therefore, a small population of cells expressing low CAG promoter activity emerge during early reprogramming.

### CAG_low_ cells enrich for reprogramming-progressive cells

To test whether the CAG_low_ cells are enriched for efficient reprogramming, we sorted the ~15% cells of the highest and lowest CAG:H2B-GFP intensity among mCherry^+^ cells and replated them separately to allow further reprogramming. As shown in Figure 2A, the CAG:H2B-GFP_low_ population displayed much higher (about 20 fold) efficiency as compared to the CAG:H2B-GFP_hi_ cells, assessed by alkaline phostphase staining. To confirm the derived colonies are mature iPSCs, we stained them for Nanog, a more stringent pluripotency marker. As expected, most colonies arose from CAG:H2B-GFP_low_ cells were Nanog-positive (Figure 2B). The expression of additional core pluripotency genes, *Oct4*, *Nanog*, *Sox2* and *Esrrb*, at similar levels as that in mESCs, further confirmed their pluripotent nature (Figure 2C). Importantly, prospective isolation of CAG:GFP_low_ cells similarly enriched for high reprogramming activity (Figure 2D). The consistent enrichment of reprogramming efficiency among the CAG_low_ cells, by two independent transgenic lines, demonstrate that the transgenic CAG promoter could reveal the reprogramming progressive cells. Therefore, low CAG promoter activity can be used to isolate early reprogramming cells capable to progress toward mature pluripotency.

**Figure 2.**
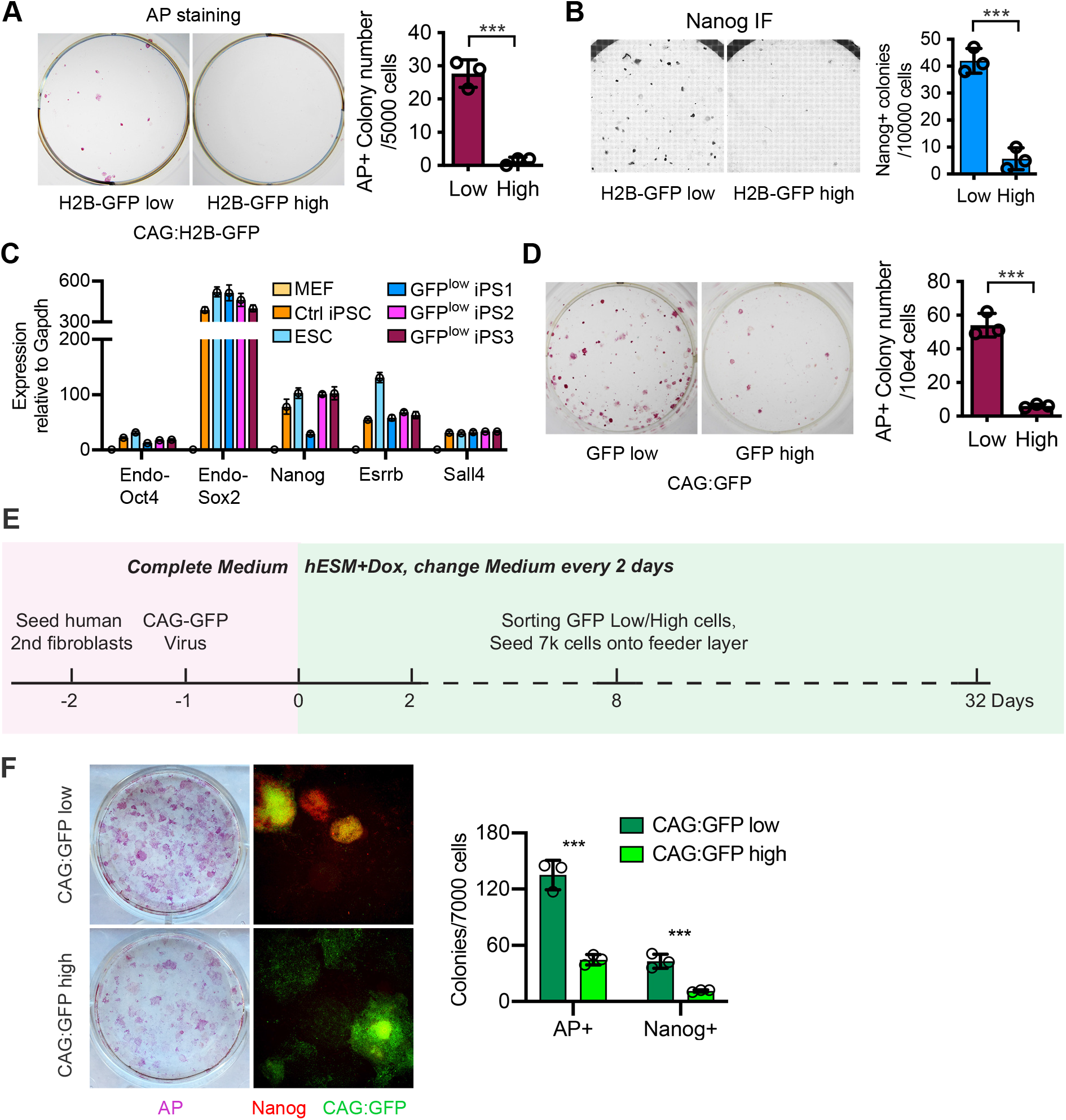
Low CAG promoter activity in reprogramming fibroblasts enriches for reprogramming progressive cells. (A) AP staining of iPS colonies at reprogramming day 12. 15% of the highest or lowest CAG:H2B-GFP+ cells were sorted from the mCherry^+^ cells on day 4 and replated on feeder cells to allow further reprogramming. Reprogramming efficiency is quantified on the right. ***: *P* < 0.001. (B) Immunostaining of iPS colonies at reprogramming day 12 for Nanog. 15% of the highest or lowest H2B-GFP+ cells were sorted from the mCherry^+^ cells on day 4 and replated on feeder cells to allow further reprogramming. Reprogramming efficiency is quantified on the right. ***: *P* < 0.001. (C) Real-time PCR analysis of core pluripotent gene expression in MEFs, control iPSCs, ESCs, and iPS colonies derived from H2B-GFP_low_ cells sorted at reprogramming day 4. Expression in MEFs is set to 1. (D) AP staining of iPS colonies at reprogramming day 12. 15% highest or lowest CAG:GFP+ cells were sorted from the OSKMmCherry^+^ population on day 6 and replated on feeder cells to allow further reprogramming. Colonies were scored and quantified on the right. ***: *P* < 0.001. (E) Schematics of reprogramming timeline using the secondary human fibroblast. (F) AP staining and Nanog immunostaining of colonies at reprogramming day 32. The numbers of AP+ or Nanog+ colonies arising from CAG:GFP-high and CAG:GFP-low cells are shown on the right. **: *P* < 0.01; ***: *P* < 0.001.

To test whether low CAG promoter activity identifies reprogramming progressive human cells, we used the secondary human fibroblasts (Hockemeyer et al., 2008;Maherali et al., 2008). These human fibroblasts contain the dox-inducible transcription factors *OCT4*, *SOX2*, *KLF4*, and *C-MYC*. We transduced the cells with CAG:GFP in a lentiviral vector, and added Dox. On day 7, the brightest and dimmest 10% CAG:GFP+ cells were FACS sorted and replated for further reprogramming (Figure 2E). Similar to the results in mouse cells, CAG:GFP_low_ cells gave rise to significantly more AP+ colonies than CAG:GFP_high_ cells. Immunostaining for Nanog confirmed positivity in many of these colonies, indicative of their mature pluripotency (Figure 2F). Taken together, our data reveal that the activity of the synthetic CAG promoter, when expressed as a transgene, has a surprising utility in enriching for reprogramming progressive cells in both human and mouse.

### CAG_low_ cells largely overlap with the fast cycling cells

Because we previously identified that a minor population of cells that have achieved sufficient cell cycle acceleration also reprogram with much higher efficiency (Guo et al., 2014), we examined the relationship between the CAG:H2B-GFP_low_ cells and the ultrafast cycling cells. Using a violet cell proliferation indicator dye (inherited by the two daughter cells of each cell division), we obtained simultaneous measurements of the CAG:H2B-GFP signal and the proliferation dye intensity. Cells with low proliferation dye intensity indicate fast cycling speed, since the dye is diluted more during the same chase period. We observed a strong correlation between the dye_low_ cells and the CAG:H2B-GFP_low_ cells, indicating that CAG:H2B-GFP_low_ cells largely overlap with the fast cycling cells (Figures 3A-B), accounting for the high reprogramming efficiency observed for both isolation approaches (Guo et al., 2014).

**Figure 3.**
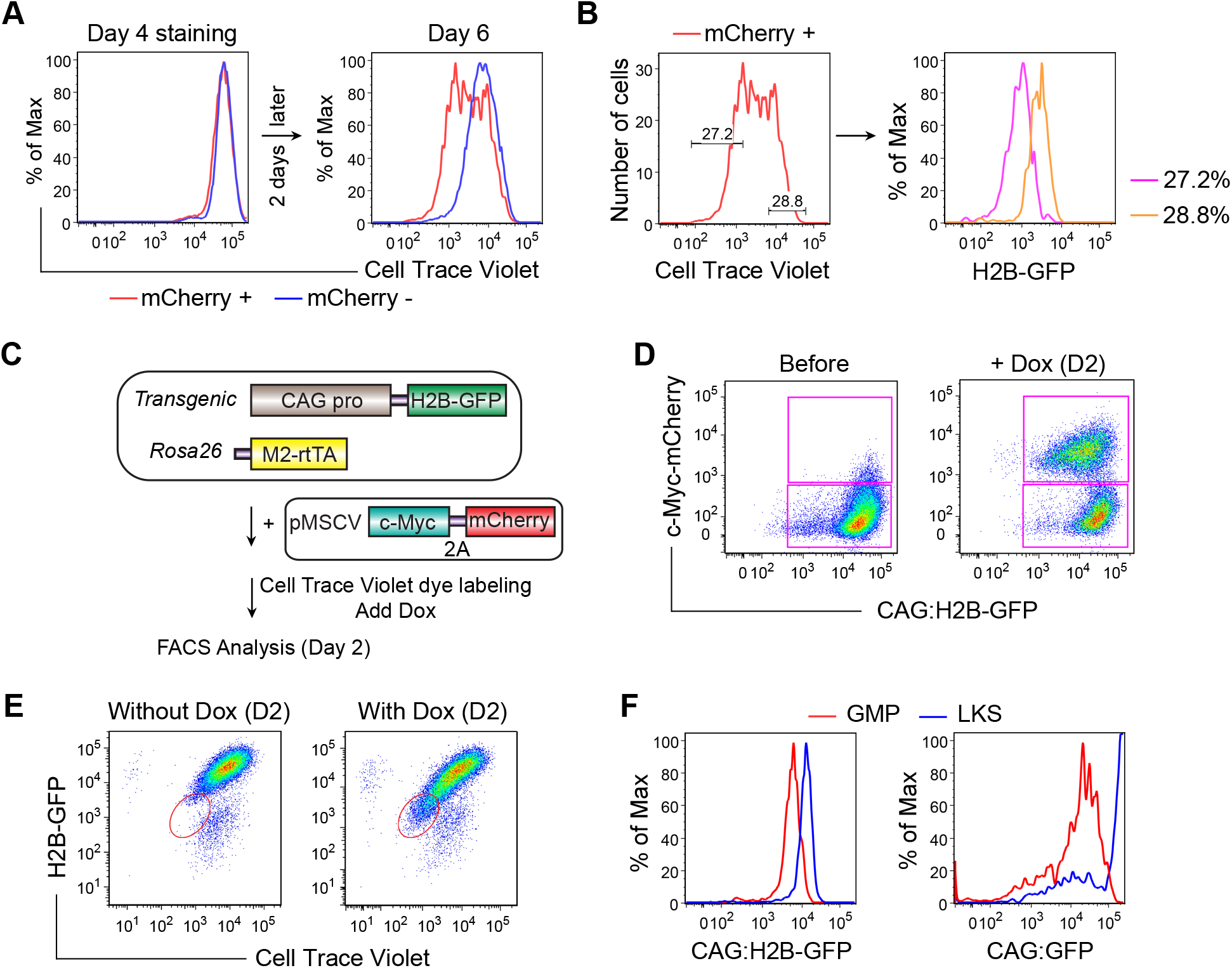
Reprogramming cells with low CAG promoter activity are fast cycling cells. (A) FACS analysis of CAG:H2B-GFP fibroblasts during reprogramming, which was performed as shown in Figure 1A. Cells were stained with Cell Trace Violet dye at day 4, replated and analyzed at day 6. mCherry^+^ and mCherry– cells indicate OSKM-expressing and non-expressing cells, respectively. (B) mCherry^+^ cells shown in (A) are gated according to the violet dye intensity, and the populations are replotted based on H2B-GFP intensity. Dye_low_ cells display low H2B-GFP intensity. (C) Schematics of the experimental design on the correlation between CAG promoter activity and c-Myc-driven cell cycle acceleration. CAG:H2B-GFP fibroblasts were transduced with inducible c-Myc-2A-mCherry. Cells were labeled with Cell Trace Violet dye and induced for c-Myc expression thereafter. (D) FACS analysis of CAG:H2B-GFP fibroblasts shown in (C), before and after induction for c-Myc expression for 2 days. Note that most of the H2B-GFP_low_ cells are mCherry^+^. (E) FACS analysis of CAG:H2B-GFP fibroblasts treated as shown in (C), without or with Dox induction for 2 days. Dye_low_ cells (oval gate) correlate with H2B-GFP_low_ cells. (F) FACS plot of fresh LKS cells and GMPs from the bone marrow of CAG:H2B-GFP and CAG:GFP transgenic mice.

To examine whether low CAG promoter activity arose when cell cycle becomes fast, we accelerated the cell cycle of CAG:H2B-GFP fibroblasts by overexpressing Dox-inducible c-Myc alone, using a vector that also encodes mCherry (Figures 3C-D). The same tight correspondence between dye retention and CAG activity was similarly observed, with dye_low_ cells displaying low CAG:H2B-GFP intensity (Figure 3E). These data suggest CAG_low_ cells could arise consequent to cell cycle acceleration, such as when c-Myc is overexpressed.

To test whether CAG_low_ cells exist in normal somatic cells of distinct cycling behavior, we examined the CAG activity in closely related hematopoietic progenitors. As shown in Figure 3F, the largely quiescent hematopoietic stem cells (Lineage– Kit^+^ Sca^+^, LKS) display higher CAG activity than the fast cycling granulocyte macrophage progenitors (GMPs), in both CAG:H2B-GFP and CAG:GFP mice. Consistent with the findings in MEFs, the CAG_low_ cells also correspond to those with high reprogramming efficiency, as GMPs reprogram more efficiently than LKS cells (Eminli et al., 2009; Guo et al., 2014). Taken together, these data demonstrate that the CAG_low_ cells are primarily the same cells as fast cycling cells, reinforcing the notion that low CAG promoter activity enrich for reprogramming progressive cells.

### CAG_low_ cells are small in size with reduced SRF target genes

To explore whether low CAG promoter activity is related to the reduced MKL1/SRF signaling pathway, we examined cell size/morphology changes in live cell imaging and tracking of cells undergoing reprogramming (Guo et al., 2014; Megyola et al., 2013), since cell size/morphology is largely determined by actin cytoskeletal genes, which are MKL1/SRF targets. We found that the reprogramming mCherry+/CAG:H2B-GFP_low_ cells formed clusters shortly after adding Dox (2 days) (Figure 1B), and these cell clusters contained cells of much smaller size. Over longer time in Dox, the clusters increased in size while individual cells within the clusters became even smaller (Figures 1B and 4A). This is consistent with earlier reports that successful reprogramming originated from cells of small sizes (Smith et al., 2010). Indeed, when we FACS sorted cells based on forward scatter alone, a crude measure of cell size, and compared their reprogramming efficiency, smaller cells indeed displayed higher reprogramming activity (Figures 4B-C). Such observation is consistent with our previous finding that reprogramming requires reduced MKL1/SRF activity (Hu et al., 2019).

**Figure 4.**
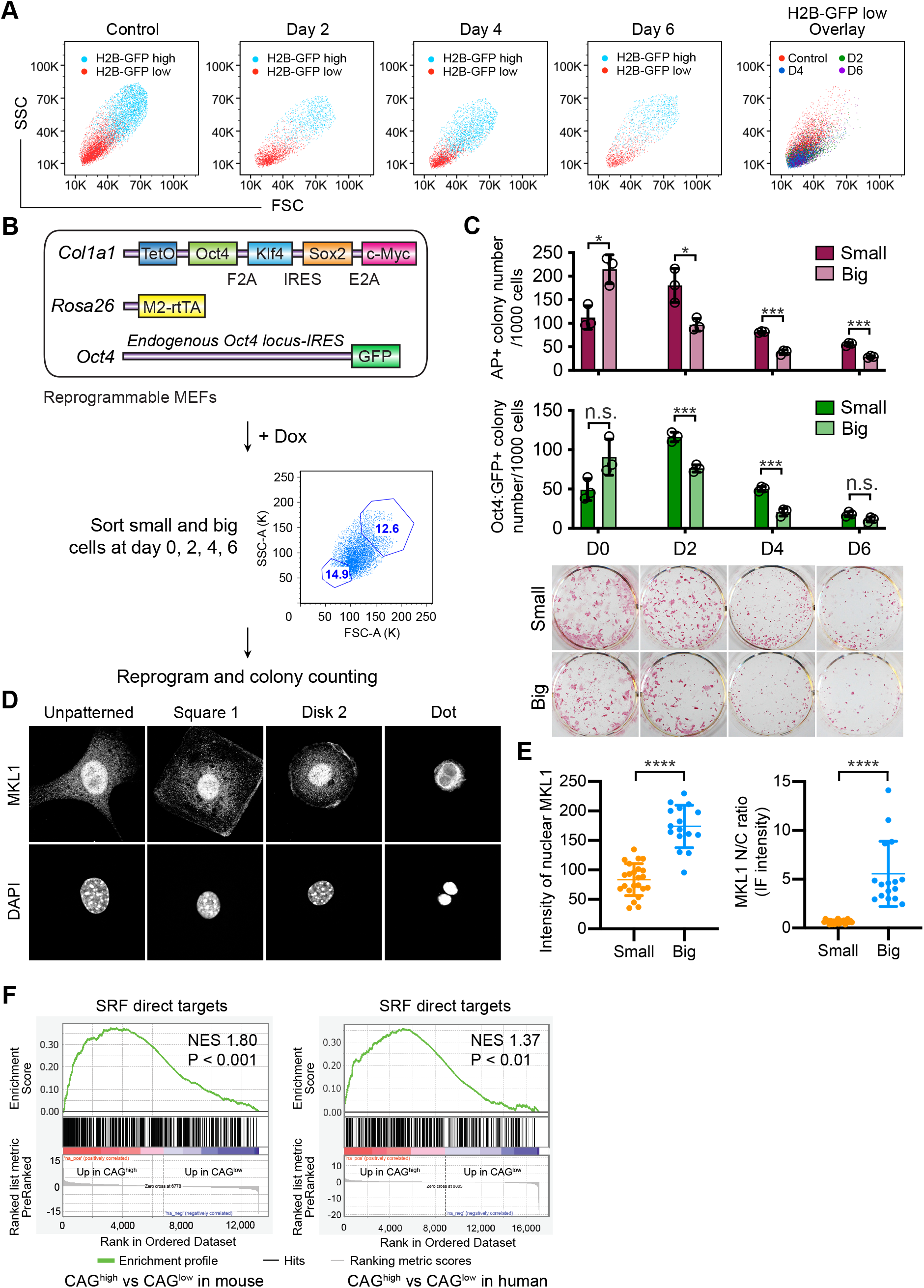
Reprogramming cells with low CAG promoter activity show reduced cell size and low MKL1/SRF activity. (A) FACS plot of H2B-GFP low and high cells at reprogramming day 0, 2, 4 and 6. H2B-GFP_low_ cells decrease in size, as indicated by decreased forward scatter (FSC) during reprogramming. (B) Design of reprogramming experiments after selecting cells based on size. Reprogrammable cells were induced for reprogramming, sorted on FSC and SSC at day 0, 2, 4, and 6, and replated on feeder cells to allow for further reprogramming. (C) Quantification of AP+ and Oct4:GFP+ colonies derived as shown in (B). Reprogramming cells with smaller size enrich for reprogramming progressive cells. *: *P* < 0.05; ***: *P* < 0.001; n.s.: non-significant. (D) Confocal images of fibroblasts grown on micropatterned surface, immunostained with MKL1 antibody, or stained with DAPI to reveal the nuclei. Micropatterned surfaces include square, disk and dot shape. (E) Quantification of MKL1 intensity in micropatterned fibroblasts. Small indicates cells patterned with dot shape, big indicates cells patterned with square and disk shape. ****: *P* < 0.0001. (F) Gene set enrichment analysis (GSEA) of differentially expressed genes between CAG_high_ and CAG_low_ cells at reprogramming day 4 (mouse) and day 7 (human). SRF target genes are enriched in the upregulated DEGs between CAG_high_ and CAG_low_ cells of the same species.

To directly monitor whether MKL1/SRF activity is modulated by cell size, we plated MEFs on micro-patterned surfaces of varying sizes, and determined endogenous MKL1 subcellular localization by immunostaining, as transcriptionally active MKL1 localizes inside the nucleus (Figure 4D). Strikingly, larger cells showed higher MKL1 nuclear to cytoplasmic ratio, while small cells had significantly decreased nuclear MKL1 (Figures 4D-E). Although the degree of enrichment for reprogramming activity was less than that achieved by dye dilution (more than 1,000 fold (Guo et al., 2014)) or CAG activity (Figure 2 above), the fact that altering just a single parameter, i.e. cell size, could change MKL1 localization strongly support that mCherry+/CAG:H2B-GFP_low_ cell clusters expressed reduced MKL1/SRF activity as they become smaller. Cell size reduction supports low MKL1/SRF activity, since this change involves dramatic reduction and rearrangement of actin cytoskeleton. The different extent of enrichment for reprogramming activity suggests that fast cell cycle likely leads to additional molecular changes besides decreasing cell sizes.

To reveal the molecular differences between the CAG_low_ and CAG_hi_ cells, conserved between mouse and human, we performed RNA-seq analyses on CAG:H2B-GFP MEFs and human secondary fibroblasts which were FACS sorted based on CAG activity on reprogramming day 4 and 7, respectively. Larger transcriptomic changes were detected between the reprogramming MEFs as compared to the reprogramming human fibroblasts, likely due to secondary nature of the human fibroblasts (Figure S1). Strikingly, however, the differentially expressed genes (DEGs) between the CAG_hi_ and CAG_low_ cells of both species revealed significantly more SRF target genes by gene set enrichment analysis (GSEA) (Figure 4F). These results confirm that the CAG promoter activity indeed report endogenous SRF activity.

Although cells with low MKL1/SRF activity enrich for reprogramming progressive cells, we note that complete lack of MKL1 interferes with MEF proliferation, and consequently reprogramming (Figure S2A-B). This observation is consistent with the fact that SRF null embryos die at E6.0 (Schratt et al., 2001), preceding the time when MEFs are derived typically on E13.5-E14.5. The difference in the severity between MKL1-null and SRF-null embryos may be related to the functional redundancy provided by MKL2 (Smith et al., 2012), as a dual targeting shRNA against both MKL1 and MKL2 ablated all AP+ colonies accompanied by reduced cell numbers (Figure S2C). Therefore, CAG_low_ instead of CAG negative cells, enrich for reprogramming progressive cells.

## Discussion

We describe a convenient approach to isolate and enrich for reprogramming progressive cells from multiple somatic cell types, of both mouse and human origin. Based on the activity of a widely used synthetic promoter, CAG, significant enrichment of reprogramming efficiency can be achieved. Specifically, cells expressing low CAG promoter activity are small in size and share identity with the previously identified fast cycling cells. Cells bearing both traits display high reprogramming efficiency. The mechanisms why CAG_low_ cells reprogram more efficiently is related to their reduced MKL1/SRF activity, as CAG_low_ cells express many SRF target genes at reduced levels. This enrichment approach is easy to implement and should help the further mechanistic studies.

We clarify the relationship between the small cells and fast cycling cells: they are essentially the same entity. Cells of small size and rapid cell cycle were reported in separate observations, both of which enrich for reprogramming cells. Specifically, early reprogramming is accompanied by dramatic cell cycle acceleration (Guo et al., 2014), and tracking pluripotency induction from somatic cells by live cell imaging have revealed that the privileged cells undergo ultrafast cell cycle of 8 hours/cycle. Independent imaging studies also revealed that successful reprogramming from fibroblasts invariably originate from cells of small sizes (Smith et al., 2010), an observation confirmed by other approaches (Wu et al., 2015). As c-Myc alone was sufficient to induce the appearance of CAG_low_ cells, and c-Myc is a potent cell cycle driver, we suggest that CAG_low_ cells arise consequent to cell cycle acceleration.

Heterogeneity, or stochasticity, in reprogramming efficiency could have many underlying reasons, including complex crosstalks with other cells. For example, reprogramming could be influenced by non-cell autonomous signals, such as IL-6 secreted by adjacent senescence cells (Brady et al., 2013; Mosteiro et al., 2016). Variability in inflammatory cytokine production (Mahmoudi et al., 2019) and cellular competition (Shakiba et al., 2019) have been recently reported to mediate different reprogramming behaviors. Furthermore, we recently described a non-cell-autonomous mode of regulation of reprogramming by the transcription co-activator YAP via one of the secreted matricellular protein CYR61 (Hartman A.S. Scalf S.M., 2018). Therefore, it could be difficult to reach absolute purification for reprogramming progressive cells. The extend of enrichment achievable by selecting for CAG_low_ cells is superior or comparable to many previously reported approaches.

## Acknowledgement

We thank Alejandro De Los Angeles for kindly providing us with the human secondary fibroblasts.

## Materials and Methods

### Mice and cells

All mouse work was approved by the Institutional Animal Care and Use Committee of Yale University. The reprogrammable mouse (*R26_rtTA_;Col1a1_4F2A_*) (Carey et al., 2010) (stock# 011004), CAG:*H2B-GFP* transgenic line (stock# 006069) and CAG:EGFP line (stock# 003291) were purchased from the Jackson Laboratory. The reprogrammable mice with reporter (*R26_rtTA_;Col1a1_4F2A_;Oct4_GFP_*) were derived by crossing reprogrammable mice with *Oct4:GFP* mice. MKL1 knockout mouse line has been described before (Smith et al., 2012). MEFs with different genetic background were all derived from E13.5 embryos. Human secondary fibroblasts were generated as previously described (Hockemeyer et al., 2008; Maherali et al., 2008) and obtained from Alejandro De Los Angeles as a gift. Granulocyte-Monocyte progenitors (GMPs) were collected from the bone marrow of mice with corresponding genotypes with the immune-phenotype of Lin-Sca-Kit+CD34+CD16/32+ as we have described before (Guo et al., 2014; Megyola et al.,2013).

### Cell culture

MEFs, 293T and human secondary fibroblasts cells were cultured in DMEM (Gibco, 11995) supplemented with 10% heat-inactivated FBS (Gibco) and 1× Penicillin-Streptomycin-Glutamine (PSG, Gibco). Mouse iPS cells and ES cells were cultured in mESC medium defined as DMEM supplemented with 15% FBS (Hyclone), 0.1 mM nonessential amino acid (NEAA, Gibco), 1 × PSG, 0.1 mM β-mercaptoethanol, and 1000 U/mL murine leukemia inhibitory factor (LIF, Millipore). Human iPS cells were cultured in mTeSRTM1 complete kit (85850). Feeder cells were obtained by irradiating P5-P6 MEFs. Mature iPS cells and ES cells were maintained on feeder layers.

### iPSC induction

For reprogramming involving Dox inducible OSKM-cassette or reprogrammable MEFs, reprogramming was induced by adding Doxycycline (Dox) to the ESC culture medium with a final concentration of 2 μg/mL. Viruses were generally transduced or cotransduced one day before initiation of reprogramming. Medium was changed every other day for reprogramming experiments. For mouse reprogramming, vitamin C was added to the reprogramming culture after replating.

### Constructs

The reprogramming factors Oct4, Sox2, Klf4 and c-Myc, and the reporter mCherry are fused with 2A, and cloned into the pFUW lentiviral backbone with an inducible promoter TetO. MKL1/2 shRNA construct was obtained from Addgene (Lee et al., 2010). CAG:GFP construct was obtained from Cellomics Technology (PLV10057). Inducible c-Myc-mCherry construct was generated by cloning the c-Myc into pFUW lentiviral backbone with an inducible promoter TetO, and a 2A-mCherry reporter.

### RNA extraction, Reverse Transcription and qPCR

Total RNA was extracted with Trizol®reagent (Invitrogen) and reverse transcribed with the SuperScript®III First-Strand Synthesis System according to manufacturer’s instructions (Invitrogen). Quantitative real-time PCR was performed using the iQ™ SYBR®Green Supermix (Bio-Rad) on a Bio-Rad CFX96.

### RNA-seq and data analysis

RNA-seq libraries were prepared with TruSeq Stranded mRNA Library Prep Kit (Illumina, RS-122-2101) following the manufacturer’s instructions. High-throughput sequencing was performed with the Illumina HiSeq 4000 Sequencing System. For data analysis, the RNA-seq reads were mapped to mouse genome (mm10) or human genome (hg38) with TopHat2. Gene abundance was calculated using cuffnorm, which outputs read counts and the number of mapped fragments per kilobase of exon per million fragments (RPKM). Genes with RPKM ≥1 in two or more samples were selected for further analysis. Differentially expressed genes were identified by Cuffdiff followed by cutting off with FDR-adjusted P value < 0.05 and fold change > 2. GSEA was performed using GSEA software (http://software.broadinstitute.org/gsea/) (Subramanian et al., 2005) with default parameters. SRF directed targets were extracted from <u>Esnault</u> et al (Esnault et al., 2014).

### FACS analysis and sorting

Cells were analyzed on BD LSRII, and sorted on BD Aria. GMPs are sorted as previously described (Guo et al., 2010).

### AP staining

AP staining was performed using the AP staining kit from StemGent (00-0055).

### Western Blot Analysis

Cell lysates were harvested by directly lysing the cells with 2 × sampling buffer (Bio-Rad). Proteins were separated by SDS-PAGE, transferred onto nitrocellulose membranes (Bio-Rad). The membranes were blocked with 5% nonfat dry milk in TBS-Tween (TBST) for 1 hour, incubated with primary antibodies overnight at 4 °C, followed by incubation with horseradish-peroxidase-conjugated secondary antibodies for 1 hour, and illuminated by enhanced chemiluminescence (ECL). Quantification of band intensity was done in Image J. β-actin antibody: Abcam, Ab20272 (1:10,000); β-tubulin antibody: Abcam, Ab6046 (1:5,000); H2B antibody: Cell Signaling, 8135 (1:2,000).

### Immunostaining

Cells or reprogramming colonies were fixed with 4% paraformaldehyde (PFA) at room temperature (RT) for 20 min, and then permeabilized with 0.5% Triton X-100 in DPBS at RT for 30 min. Samples were blocked with blocking buffer (5% normal goat serum (NGS), 0.3% Triton X-100 in DPBS) at RT for 1 hour. Primary antibody incubation was performed at 4°C overnight. Antibody was diluted with antibody buffer (1% BSA, 0.3% Triton X-100 in DPBS). After washing, samples were then incubated with secondary antibody prepared with antibody buffer at RT for 1 hour. For micropatterning, images were taken under a confocal microscope. For iPS colony quantification, images were taken with the live cell imaging system (Molecular Devices). Nanog antibody: Cell Signaling, 4903 (1:200); MKL1 antibody: a gift from Topher Carroll.

### CFSE/Cell Trace Violet dye staining

CFSE/Cell Trace Violet dye staining was performed according to the manufacturer’s instructions. Briefly, cells were trypsinized, washed, and resuspended with DPBS in 1 million cells/mL concentration. Cells were then incubated with CFSE/Cell Trace Violet dye at 37 °C for 20 min, and supplied with 5 times the volume of complete culture medium, followed by incubation at 37 °C for 5 min. Cells were centrifuged and resuspended in prewarmed culture medium with incubation for more than 10 min. And then, they were subject for either flow analysis or replating for further growth.

## Supplementary Figures

**Figure S1.**
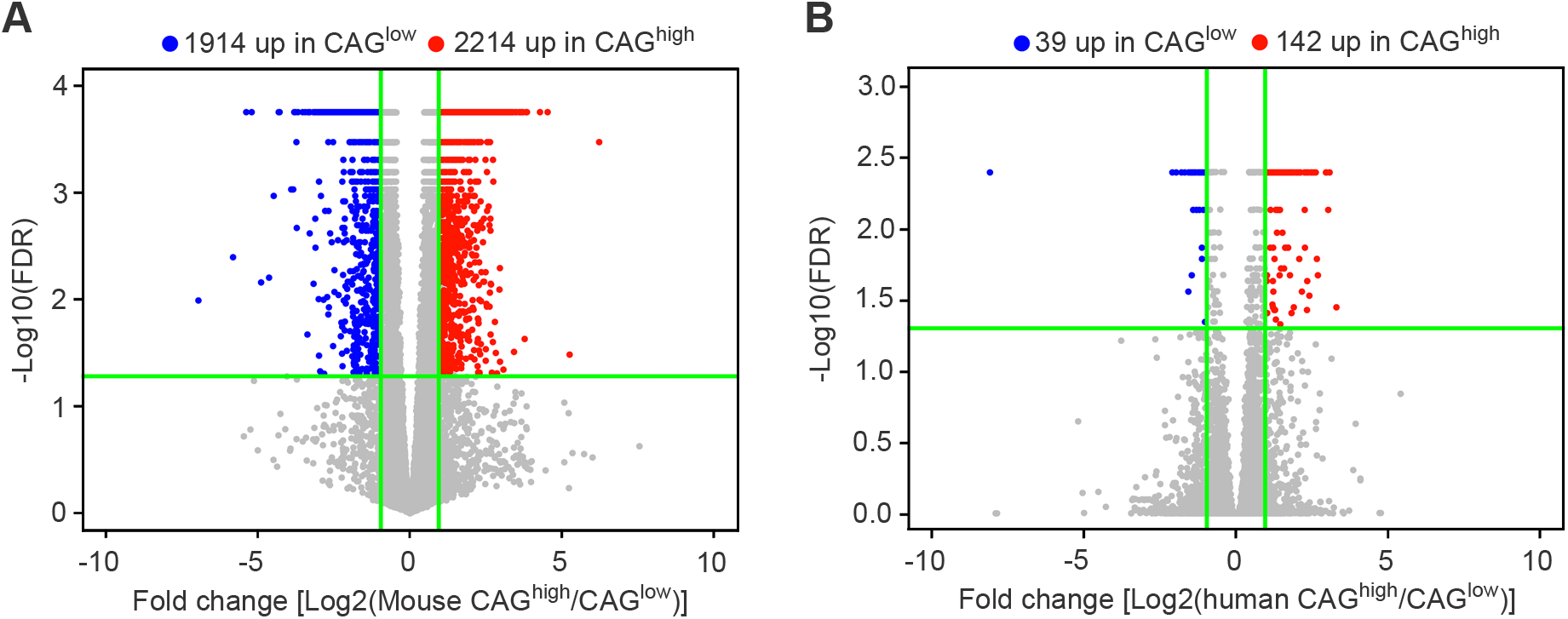
CAG_Hi_ and CAG_Low_ cells display different gene expression patterns. (A) Defferentially expressed genes in CAG_Hi_ and CAG_Low_ cells derived from mouse fibroblasts. (B) Defferentially expressed genes in CAG_Hi_ and CAG_Low_ cells derived from human fibroblasts.

**Figure S2.**
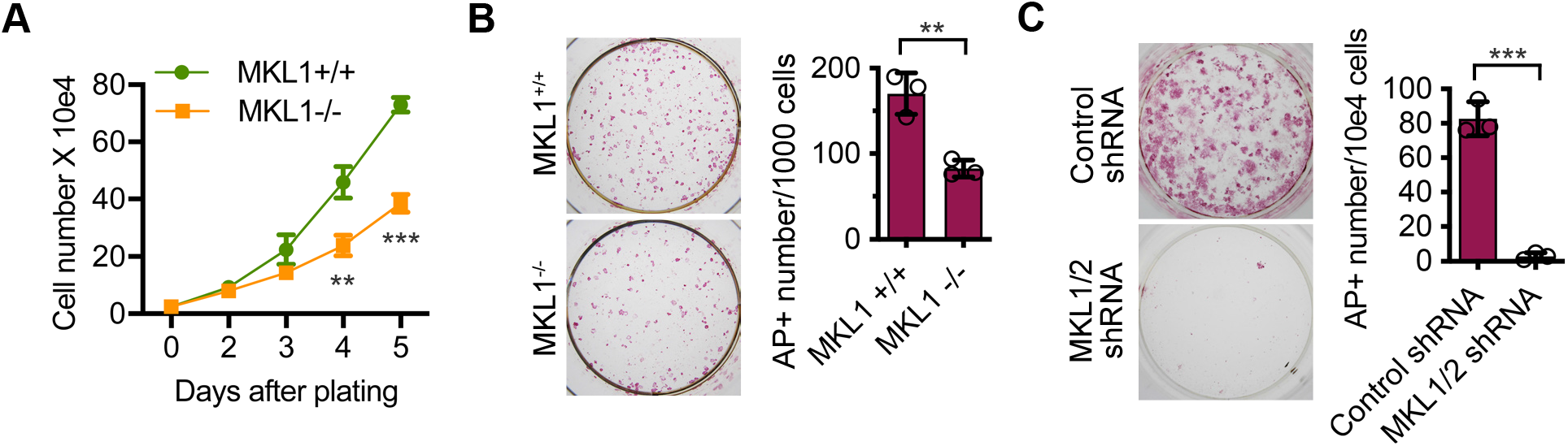
Complete MKL1 loss-of-function impairs fibroblast proliferation and inhibits reprogramming. (A) Population doubling time of primary MKL1_+/+_ and MKL1_−/−_ MEFs. **: *P* < 0.01; ***: *P* < 0.001. (B) AP staining and quantification of colonies induced from MKL1_+/+_ and MKL1_−/−_ MEFs at reprogramming day 10. **: *P* < 0.01. (C) AP staining and quantification of colonies transduced with control shRNA or MKL1/2 shRNA at reprogramming day 10. ***: *P* < 0.001.

